# SAFE-OPT: A Bayesian optimization algorithm for learning optimal deep brain stimulation parameters with safety constraints

**DOI:** 10.1101/2024.02.13.580142

**Authors:** Eric R. Cole, Mark J. Connolly, Mihir Ghetiya, Mohammad E. S. Sendi, Adam Kashlan, Thomas E. Eggers, Robert E. Gross

**Affiliations:** Wallace H. Coulter Department of Biomedical Engineering, Georgia Institute of Technology and Emory University, Atlanta, GA, 30332, USA; Department of Neurosurgery, Emory University School of Medicine, Atlanta, GA, 30322, USA; Emory National Primate Research Center, Atlanta, GA, 30322, USA; Emory College of Arts and Sciences, Emory University, Atlanta, GA, 30322, USA; College of Sciences, Georgia Institute of Technology, Atlanta, GA, 30322, USA; Department of Neurosurgery, Robert Wood Johnson Medical School, Rutgers University, New Brunswick, NJ, 08901, USA

**Author notes:** **Corresponding Author**: Robert E. Gross, MD, PhD, Department of Neurosurgery, 10 Plum Street, New Brunswick, NJ 08901. Equal contribution (Eric R. Cole, Mark J. Connolly).

**Keywords:** Neuromodulation, optimization, hippocampus, data-driven, real-time

## Abstract

To treat neurological and psychiatric diseases with deep brain stimulation, a trained clinician must select parameters for each patient by monitoring their symptoms and side-effects in a months-long trial-and-error process, delaying optimal clinical outcomes. Bayesian optimization has been proposed as an efficient method to quickly and automatically search for optimal parameters. However, conventional Bayesian optimization does not account for patient safety and could trigger unwanted or dangerous side-effects. In this study we develop SAFE-OPT, a Bayesian optimization algorithm designed to learn subject-specific safety constraints to avoid potentially harmful stimulation settings during optimization. We prototype and validate SAFE-OPT using a rodent multielectrode stimulation paradigm which causes subject-specific performance deficits in a spatial memory task. We first use data from an initial cohort of subjects to build a simulation where we design the best SAFE-OPT configuration for safe and accurate searching *in silico*. We then deploy both SAFE-OPT and conventional Bayesian optimization in new subjects *in vivo*, showing that SAFE-OPT can find an optimally high stimulation amplitude that does not harm task performance with comparable sample efficiency to Bayesian optimization and without selecting amplitude values that exceed the subject’s safety threshold. The incorporation of safety constraints will provide a key step for adopting Bayesian optimization in real-world applications of deep brain stimulation.

## 1. Introduction

Deep brain stimulation (DBS) and other neuromodulation approaches are increasingly being used for treating neurological and psychiatric disorders such as treatment-resistant epilepsy, depression, and Parkinson’s disease.^1–5^ A critical step in delivering effective therapy is selecting the correct stimulation setting – a combination of stimulation parameters, such as the amplitude, frequency, and spatial configuration of electric current delivered to the brain – for each individual patient.^6, 7^ Due to variability in electrode placement, patient anatomy, and physiology, the optimal stimulation setting for a given patient is not known *a priori*. Instead, it must be identified through a tedious trial-and-error process whereby a clinician applies the stimulation setting and observes the patient until the effect can be measured, and then repeats this process until achieving the desired effect. This practice poses several challenges, because it can take a considerable amount of time to measure the effect of a given stimulation setting – up to one week for depression^5^ and months or years for epilepsy.^8, 9^ Therefore optimizing even a single stimulation parameter (e.g., amplitude) can nearly become an intractable challenge.

Data-driven, particularly Bayesian, optimization has recently been proposed as an algorithmic strategy to efficiently search for the optimal stimulation setting for each individual patient.^10–14^ These approaches have been shown to be robust to noise, are more efficient than more naïve grid or random search approaches, and are capable of navigating high-dimensional parameter spaces.^15–17^ Data-driven optimization has the potential to automate the process of selecting stimulation settings, paving the way for continually learning the optimal stimulation setting for an individual patient without ongoing supervision from clinicians.

For optimization to be used in clinical practice, it needs to account for patient-specific safety considerations. While DBS is generally well-tolerated in patients, regions of the parameter space that are optimal in some subjects can produce uncomfortable and possibly dangerous side-effects in other subjects. These side-effects include muscle contractions, voice changes, deleterious mood and cognitive changes, memory impairments, seizures, and more.^18–23^ While side effects or returning symptoms caused by a change to the stimulation setting may be briefly tolerable under the supervision of a clinician, they could be unsafe or detrimental during daily living activities. As a consequence, data-driven methods must be designed to actively learn and avoid unsafe parameter space regions for any automated or unsupervised optimization approaches. This can be accomplished using recent work which demonstrates how Bayesian optimization can be augmented with learnable safety constraints to safely and efficiently navigate a parameter space.^24^

This study uses a previously described design framework^11^ to prototype and deploy a Bayesian optimization algorithm with learnable safety constraints, which we name SAFE-OPT (**Figure 1**). We designed, characterized, and deployed this algorithm in intact, healthy rats undergoing a spatial object recognition (SOR) memory task while receiving hippocampal electrical stimulation. The stimulation paradigm used was originally developed to reduce seizures in the rat tetanus toxin model of temporal lobe epilepsy,^25^ and was later shown to impair performance on the SOR task at sufficiently high stimulation amplitudes that varied between subjects.^26^ Using the effects of multielectrode stimulation on SOR performance as a model system, we designed and validated a proof-of-concept algorithm that can efficiently find the subject-specific optimal stimulation amplitude and safety threshold while carefully avoiding regions of the parameter space that could cause memory deficits.

**Figure 1:**
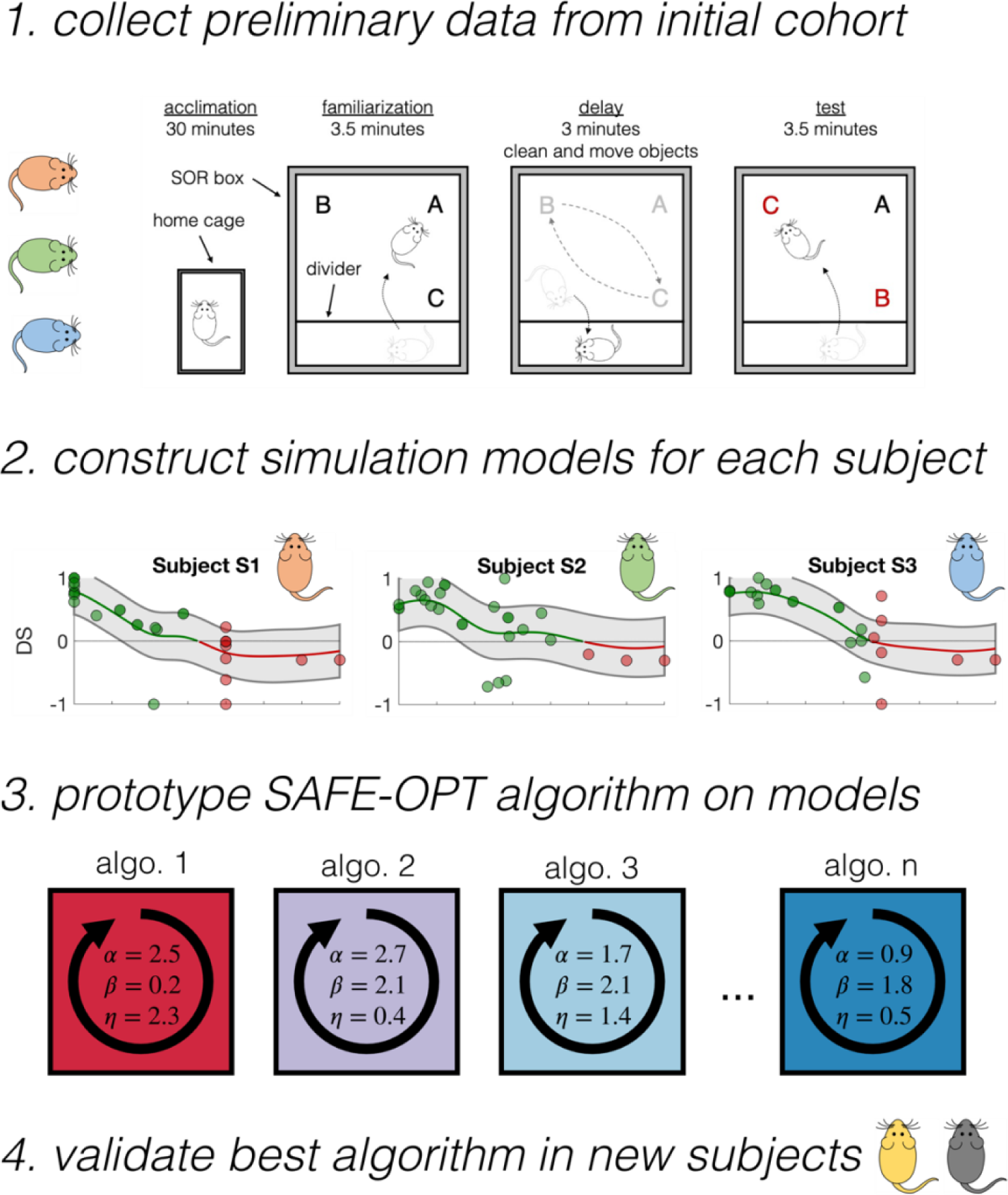
Framework for designing SAFE-OPT. A four-phase methodology was used to develop and test the SAFE-OPT algorithm. 1) Data Collection: In a cohort of multiple subjects, many different stimulation settings are applied and the resultant performance on a memory task is measured for each setting. 2) Modeling: The data from each subject is used to fit a regression model that predicts both the mean and variability of the task score as a function of stimulation parameters. 3) Prototyping: Many different configurations of SAFE-OPT are deployed in a simulation using each of the subject-specific models, and are evaluated for speed, safety, and consistency in finding the best stimulation setting. 4) *In vivo* validation: The configuration of SAFE-OPT that performed best across the simulation models is then validated *in vivo* in a new cohort of subjects.

## 2. Methods

The organization of the methods follows the four main steps in the design framework: 1) data collection, 2) modeling, 3) algorithm prototyping, and 4) *in vivo* validation (**Figure 1**). Section 2.1 describes the experimental setup and the data collected from an initial cohort of subjects. In Section 2.2, this data is used to construct high-throughput ground-truth statistical models for each individual subject. In Section 2.3, these models are used to rapidly prototype many variations of safe optimization algorithms. Finally, in Section 2.4 these algorithms are deployed *in vivo* to validate the performance of the algorithms in a prospective cohort of subjects.

### 2.1 Data Collection

#### 2.1.1 Subjects and Surgery

A total of 4 adult male Sprague-Dawley rats (2-3 month old; 250-300 g) from Charles River Laboratories (Wilmington, MA, USA) were used for the study along with data from 4 additional subjects that were collected as part of a previous study.^26^ All animals were maintained within a 12/12 light/dark cycle vivarium with unlimited access to food and water. All procedures were conducted in accordance with Emory University’s Institute for Animal Care and Use Committee.

Each subject underwent a single survival surgery as previously described.^25^ Under 1-4% isoflurane anesthesia a 16-channel multielectrode array (MEA; Tucker Davis Technologies (TDT), Alachua, FL., USA) was implanted targeting the CA3 and CA1 regions of the right hippocampus, centered at 3.50 mm posterior and 2.80 mm lateral to bregma. The MEA was lowered ventrally into the brain until single unit activity was observed from both the CA1 and CA3 regions. Five 2-mm stainless steel screws were mounted on the skull to serve as the ground and reference of the electrode and for structural support.

#### 2.1.2 Asynchronous distributed stimulation

The stimulation pattern based consisted of symmetric square wave biphasic pulses with 400 µs per phase on eight stimulating contacts (due to the electrode design, where eight channels were designed for active amplification during recording and eight served as passive channels for stimulation) (**Figure 2B**). Stimulation was applied fully asynchronously, where the stimulation waveform was equivalent across all contacts aside from the phase offset. Each contact had a pulse frequency of 7 Hz and a phase offset of 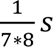. Stimulation was voltage controlled with amplitudes ranging from ±0-5 V.

**Figure 2:**
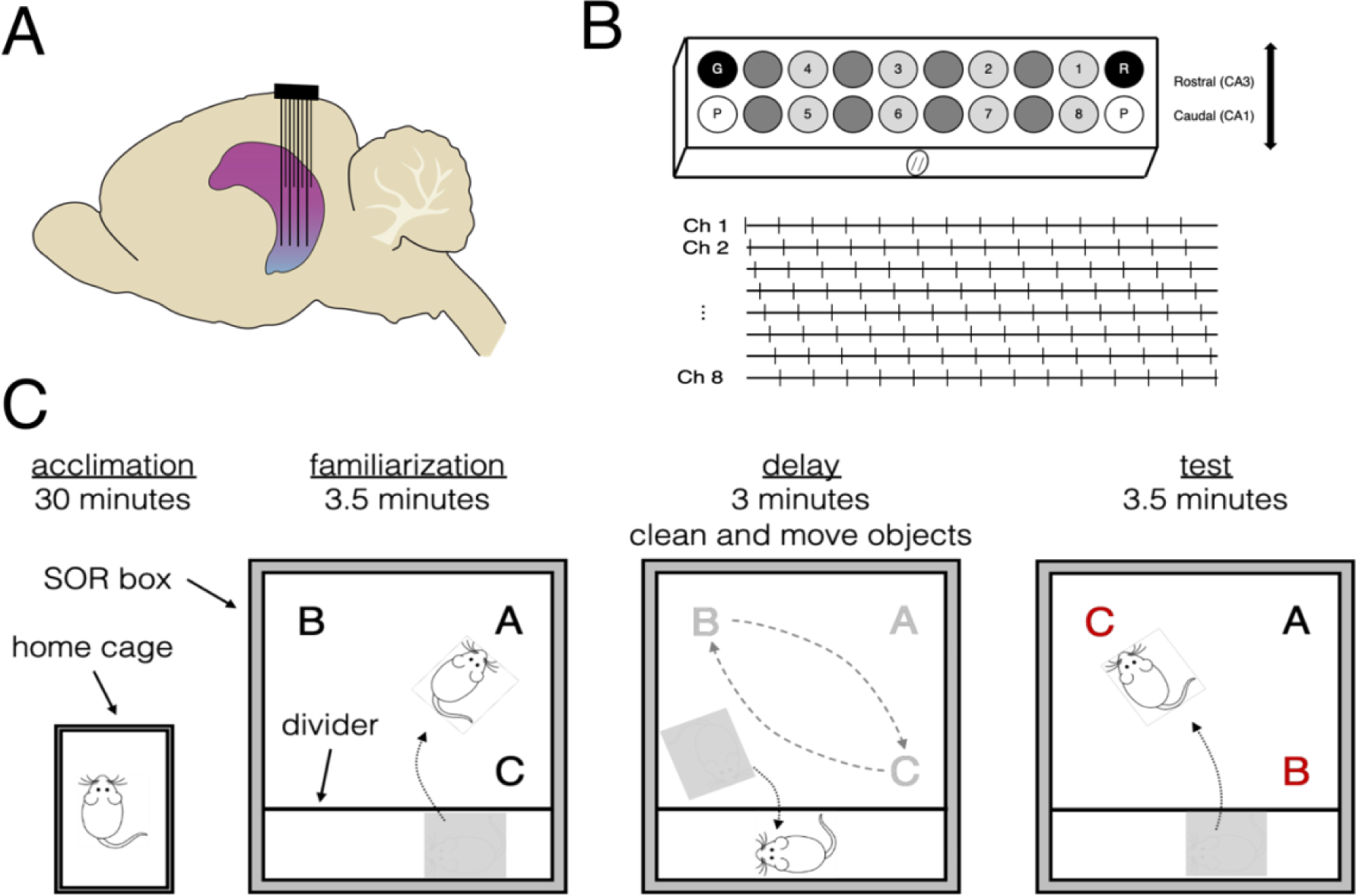
Experiment design for multielectrode stimulation during a spatial object recognition task. **A)** Staggered electrode array design with rows targeting hippocampal CA1 and CA3. **B)** Top: Electrode array channel assignment (G: ground; R: reference; P: guide pin. Stimulating channels are numbered). Bottom: the asynchronous distributed stimulation pattern is delivered in the form of a constant-frequency pulse train with a phase offset across channels. **C)** Spatial object recognition (SOR) task design. *<u>Acclimation phase</u>*: the subject acclimates to the experiment room while remaining in their home cage for 30 minutes. The subject is then connected to a headstage and placed in the SOR box, separated from the objects by an opaque removable divider. *<u>Familiarization</u> <u>phase</u>*: stimulation begins. Then after 1 minute, the divider is removed, and the subject interacts with three objects (A, B, and C). After 3.5 minutes, the subject is again separated from the field. C) *<u>Delay phase</u>*: The field and objects are cleaned, the objects are returned, and the positions of objects B and C are swapped. D) *<u>Test phase</u>*: After 3 minutes, the divider is again removed, and the subject interacts with the objects for 3.5 minutes. A discrimination score is measured using the subject’s time spent interacting with objects during the test phase. Each trial of the SOR task produces a single DS measurement.

#### 2.1.3 Spatial object recognition task

The SOR task is a well-established behavioral task to test object recognition memory in various animal species^27^ which relies on the innate preference of rats for novel objects and changes in object location. Before starting the SOR experiments, the implanted rats were habituated to a 2’x1.5’ open-field SOR box for 5 minutes per day for three days. Practice objects were placed in the SOR box on the third day to determine whether the subjects were willing to explore the objects or additional habituation was necessary. One session of the SOR task consists of four phases: acclimation, familiarization, delay, and test. (**Figure 2**). In the acclimation phase, the rat was moved to the testing room 30 minutes before the SOR task was performed. Meanwhile, the SOR box was thoroughly cleaned with 70% isopropyl alcohol to remove any residue that could prompt olfactory sensation, and three distinct objects were placed in the corners of the exploration area. At the end of the acclimation phase, the headstage was connected to the electrode on the subject, who was then moved to a region of the SOR box separated from the exploration area by an opaque divider. The headstage was connected to a commutator suspended above the SOR box and stimulation was started. After 1 minute of stimulation, the divider was temporarily removed to begin the familiarization phase. The subject was allowed to interact with the objects for 3.5 minutes, and then the divider was again used to separate the subject from the exploration area for 3 minutes. During this delay phase, the exploration area and the objects were again thoroughly cleaned with 70% isopropyl alcohol to remove any olfactory signals. The positions of two objects were interchanged and the position of the third object was kept constant (**Figure 2**). During the testing phase, the subject was returned to the exploration area and allowed to explore the objects for 3.5 minutes. The rat was then removed from the exploration area to complete the task.

To prevent bias, the orientation of the three objects was randomized between trials. A camera placed above the SOR box captured video during each familiarization and testing phase. SOR experiments were performed in a sound-isolated room with limited red light to facilitate exploration, as rodents are nocturnal.

#### 2.1.4 Discriminant Score

The discrimination score (DS) was used to quantify the animal’s memory performance. DS is defined as the difference in mean exploration time for interchanged objects and exploration time for the remaining stationary object, divided by the sum of both quantities:

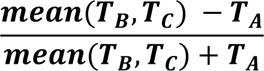

Where ***T_A_***, ***T_B_*** and ***T_C_***, represent the time the subject spent interacting with the three objects. The score ranges from −1 to +1. A score of +1 indicates complete preference for the interchanged objects, and a score of −1 indicates preference only for the stationary object. A score of 0 indicates equal preferenence for all three objects and suggests that the subject displayed no memory function, whereas a score near 1 indicates a high preference for the novel object and therefore healthy memory. SOR trials where stimulation caused an adverse reaction (such as electrographic afterdischarges, seizures, etc.) were stopped and scored with a DS of −1, indicating an unsafe effect for the given stimulation setting. Object exploration was defined by periods when the orientation of the subject’s snout was directed towards the object and animal was actively whisking indicating direct interaction. Other possible interactive behaviors, such as running around the object, playing with the object, or sitting and climbing on the object were not measured as exploration. Videos recorded during the SOR task were scored by a blinded observer using Spike2 (Cambridge Electronic Design Limited, Cambridge, England) to annotate the start and stop times of each object interaction period.

Each subject performed only one SOR task per day to prevent sequential effects of stimulation on subsequent trials or changes in performance due to fatigue. If the headstage was disconnected or the experiment was otherwise interupted during the SOR task, the individual trial was discarded and experiments were resumed the next day.

#### 2.1.5 Collected data and data augmentation

The data used in the design framework was collected from grid search and optimization experiments for 4 subjects conducted as part of a previous study^26^, as well as the 2 subjects included in the standard Bayesian optimization *in vivo* experiments in Section 3.4.1. For some subjects, the maximum amplitude tested (±5 V) did not disrupt memory, i.e., the average DS did not drop below 0. In these cases, the dataset was augmented with artificial data points at ±6 and ±7 V, to create a region of the parameter space with an average DS of 0 (visualizable in **Figure 3**).

**Figure 3:**
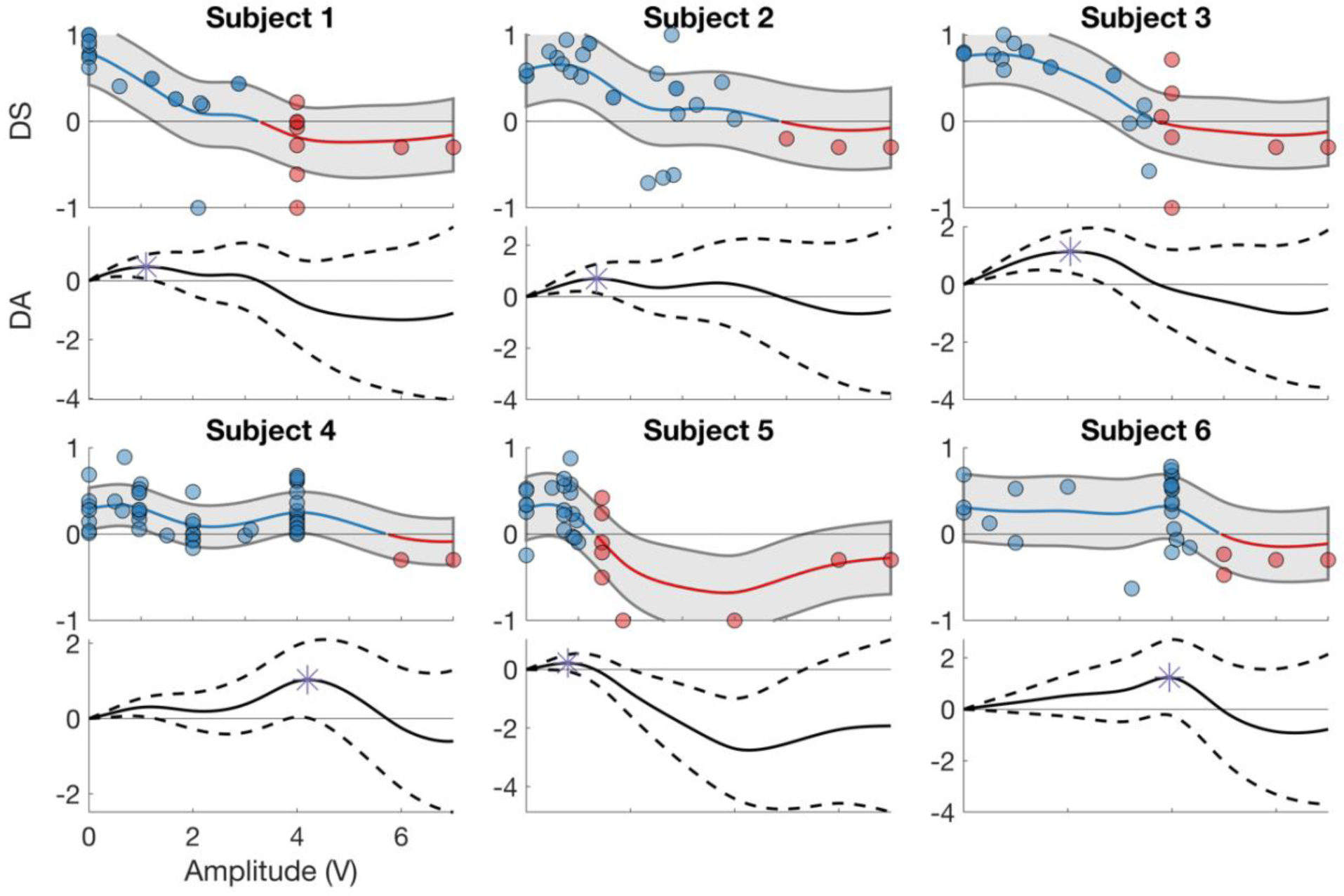
Preliminary data and ground-truth simulation models. Subject-specific simulation models mapping stimulation amplitude to the DS and the corresponding DA objective function. DS plot (top): the blue line highlights the region of the stimulation parameter space where the expectation of the DS is greater than 0. The red line indicates an expected DS less than zero. The blue and red circles show the data used to construct the ground-truth models where the expected DS is greater than and less than zero, respectively. The gray region indicates one standard deviation above and below the expectation. DA plot (bottom): solid and dashed lines indicate the expected DA based on the DS model and the standard deviation, respectively. Purple asterisk: estimated optimum (maximum DA and corresponding stimulation amplitude) of the DA objective function.

### 2.2 Ground-truth models for simulating optimization

#### 2.2.1 Gaussian process modeling

We considered the performance on the SOR task, the DS, as the safety function with a threshold of 0. The means that the goal is to optimize stimulation without testing an amplitude with an average DS of zero, indicating impaired memory. Using the data for each subject, the input (stimulation amplitude) and corresponding outputs (DS) were fit to a Gaussian process regression model as described in our prior work.^11, 28^ The Gaussian process models were composed of mean and covariance functions:

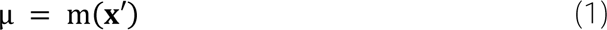

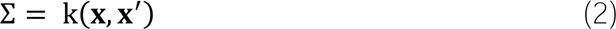

where *x* is a vector of the data used to construct the model, and *x*′ is an unlabeled sample point. The input to the model was a one-dimensional parameter space representing the stimulation amplitude, discretized to a ±0.05 V resolution. The output of the model was the DS drawn from a Gaussian distribution.

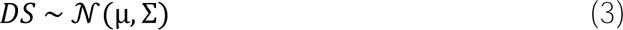

The GP was a constant mean function and a third order Matérn kernel with automatic relevance determination for the covariance:

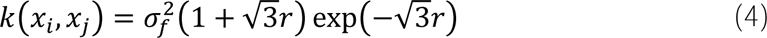

where 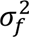 is the variance of the output and:

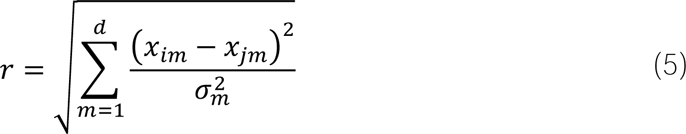

where 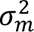 is the length scale hyperparameter which describes the variance of the stimulation amplitude parameters, *x*_*m*_. In other words, 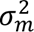 determines the extent to which the expectation can vary between adjacent stimulation amplitudes – the higher the length scale value, the more similar the expectation between neighboring amplitude values. Model hyperparameters (mean, length scale, signal noise, measurement noise) were fit by likelihood maximization. Throughout this manuscript, the term “hyperparameter” will denote adjustable parameters that determine GP model fit, whereas “configuration parameters” are other adjustable parameters determining the Bayesian optimization search strategy.

#### 2.2.2 Safety objective function: the “discrimination area”

For the objective function of optimization, we chose not to directly maximize the discrimination score or stimulation amplitude, as the optimum would simply be the edge of the safety function. Instead, we constructed an alternative objective function, the “discrimination area” (DA):

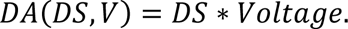

By construction, this objective function possesses several convenient properties. Specifically, when either the amplitude or DS are equal to zero, the DA is also zero. However, at a midpoint when stimulation with an amplitude greater than zero is applied and does not completely disrupt memory, the DA is positive. Similarly, because the GP features fundamental properties of differentiability and smoothness, there must exist at least one stimulation amplitude for which the derivative is zero, which based on previous data is likely a non-trivial maximum of the objective function. The objective function was modeled using a separate Gaussian process.

### 2.3 SAFE-OPT: Bayesian optimization with learnable safety constraints and hyperpriors

#### 2.3.1 Bayesian optimization

The Bayesian optimization algorithm used in this study was based on a GP model and an upper confidence bound (UCB) acquisition function.^29^ The algorithm is initialized by first applying a randomly sampled set of burn-in stimulation settings (amplitude). The objective function (DA) is calculated and the data is fit to a surrogate GP model. An acquisition function is then used to identify regions of the parameter space where the model predicts the average DA would be high (exploitation), but also has high uncertainty with the statistical potential for an even higher DA value (exploration). The UCB acquisition function is defined as:

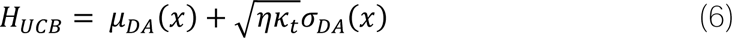

Where the configuration parameter, *η*, quantifies the tradeoff in emphasizing exploitation vs. exploration. The sequence *κ*_*t*_ was defined as:

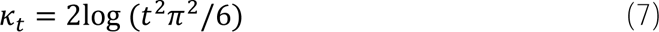

where *t* is the number of samples collected thus far during a single optimization sequence.

#### 2.3.2 SAFE-OPT: Bayesian optimization with learnable safety constraints

To implement Bayesian optimization with learnable safety constraints, we extended the UCB acquisition function with an additional safety criterion designed to avoid stimulation settings that are expected to produce a DS below zero. When selecting a stimulation setting, the surrogate DS model was queried to identify regions of the parameter space that satisfied the criteria:

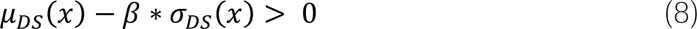

where *β* was a configurable parameter to control how carefully the algorithm would avoid potentially unsafe regions of the parameter space. A high *β* would cause the algorithm to avoid stimulation settings with even a small risk of producing a mean DS below zero. However, a *β* near or equal to zero would cause the algorithm to only avoid regions of the parameter space where it expects a mean DS of zero or lower.

#### 2.3.3 Bayesian optimization with hyperpriors

The algorithm was further extended to incorporate information from prior subjects in the form of hyperpriors, which could further improve the performance of the algorithm by improving the model fit when only a few samples have been collected.^28, 30^ When fitting the model by maximizing the likelihood over the space of hyperparameters (as defined in Section 2.2), a prior distribution was placed on the hyperparameters. This distribution was calculated based on the ground-truth models for each *other* subject. The GP kernel used in this study has hyperparameters for the mean function, the length scale for the stimulation amplitude parameter, the signal noise, and the measurement noise. For each of the hyperparameters, the mean and variance across subjects was calculated and used to define a Gaussian distribution hyperprior for that hyperparameter. When fitting the model during optimization, the Gaussian distribution hyperprior was defined for each of the hyperparameters using these statistics. A configuration parameter, *α*, was multiplied by the variance of the hyperprior, such that a small *α* would emphasize hyperparameters that are close to the hyperprior mean, while a large *α* would allow a broader range of hyperparameters to be considered. After each additional stimulation setting was tested, the model was re-fit using the hyperpriors and the entire parameter space re-estimated to compute the acquisition and safety functions from the surrogate model.

### 2.4 Simulation experiments

*In silico* experiments were performed by deploying each algorithm to search for an optimal stimulation setting using the ground-truth simulation models. To simulate an optimization trial, we extended a methodology developed in our prior work where the optimization algorithm is allowed to choose a given stimulation setting and apply it as an input to the ground-truth GP model^11^. An outcome value is randomly sampled from a Gaussian distribution using the mean and variance of the GP model at the given input value. This sample is then provided as a new data point to the algorithm, and the process iterates until convergence.

The configuration parameters for Bayesian optimization with learnable safety constraints and hyperpriors were characterized by performing a comprehensive sweep of different parameter combinations. The configuration parameters were selected from uniform distributions of η (exploration) = [0 4], β (safety) = [0 3], and α (hyperprior) = [0 3]. For simulation models corresponding to each of the six subjects, a predefined set of 30 configuration parameter combinations was evaluated for 15 trials (a single sequence where the optimization algorithm searches for an optimal stimulation setting) of 15 samples each (the number of stimulation settings tested in one trial).

### 2.5 Algorithm performance criteria

The algorithms were evaluated based on two performance criteria: final error and overshoot. Final error was calculated as the difference between the DA produced by the estimated optimal stimulation amplitude when applied to the ground-truth model and the max of the ground-truth model, representing the algorithm’s accuracy in identifying the optimal stimulation setting. The overshoot, representing the algorithm’s safety performance in navigating stimulation settings, was calculated as the difference between the maximum stimulation amplitude that was tested at any point during the optimization trial and the estimated optimal stimulation amplitude.

### 2.6 *In vivo* optimization

#### 2.6.1 Optimization phase

Each optimization trial was initialized with a set of presumed safe stimulation settings, providing the burn-in trials used by the optimization algorithm: 0, 0.5, and 1.0 V. Each stimulation setting was applied during one trial of the SOR task, the DS was scored for each trial, and the parameter and DS data were used to fit the surrogate GP model. Then the acquisition function was used to select new stimulation settings for each SOR trial, until meeting a stopping criterion (the algorithm’s estimated optimal stimulation setting remained the same for three samples, or 30 samples total – whichever came first). During optimization, data from every valid SOR trial were included, even if the subject did not interact with all objects in the test phase of the SOR task (e.g. if the subject was uninterested in the task and spent very little time exploring on a given day). Examples of invalid trials included experiments that were concluded early or where the subject did not interact with any object, producing an undefined DS value.

#### 2.7.2 Validation phase

In the validation phase, we performed 15 additional SOR trials post-optimization to compare the optimal stimulation setting estimated by the algorithm, ±0V sham stimulation, and a subject-specific control point specified as the lower of either *x*^∗^ + 1*V* or 2 · *x*^∗^ (where *x*^∗^ is the optimal stimulation setting) – an amplitude value expected to be unsafe for the given subject. The trial order was randomized between the three parameter settings. If the control point did not produce a memory deficit in a given subject, an additional control point at *x*^∗^ + 2*V* was also evaluated.

## 3. Results

We first provide a visualization of preliminary data and corresponding subject-specific GP model predictions that provide the basis for simulation. Second, we show the simulation results used to determine the optimal parameter configuration for SAFE-OPT. We then compare the results of deploying optimally designed SAFE-OPT to those of standard Bayesian optimization *in vivo*, demonstrating improved safety performance in navigating stimulation settings while maintaining comparable performance in finding the optimum.

### 3.1 Data collection

Stimulation and memory performance data were collected from six male Sprague-Dawley rats between 16-50 weeks of age. The data from each subject included the stimulation amplitude and DS for the corresponding SOR task.

### 3.2 Ground truth models

Ground-truth models were constructed using all available data for each of six subjects. The stimulation amplitudes ranged from ±0-5 V, with two additional artificial data points at ±6 and 7 V (**Figure 3**). As measured by the GP model DA predictions, the estimated optimal stimulation amplitudes ranged from ±1.1 V to ±4.0 V, with safety thresholds (i.e. the lowest stimulation amplitude where predicted DS is negative) ranging from ±1.8 V to ±4.8 V across the different subjects. All subjects demonstrated a one-sided safety boundary: all stimulation amplitudes greater than the subject-specific safety thresholds were also estimated to be unsafe, according to the expectation of each simulation model.

### 3.3 SAFE-OPT algorithm design

The SAFE-OPT algorithm configuration was determined by first quantifying the effect of configuration parameters on optimization performance across the six simulation models (**Figure 4A**). Final error (accuracy in finding the optimal amplitude) and overshoot (safety) metrics were compared for the best, average, and worst performance of each of the 30 combinations of configuration parameters across multiple subject models and multiple randomized trials of the simulation. The best and average final error values were not significantly influenced by the algorithm configuration, though there was more variability for the worst-case final error. In contrast, the overshoot was sensitive to changes in the configuration settings, specifically β (**Figure 4B**). As β is increased, the average and worst case overshoot significantly decreases (linear regression; p < 0.001), indicating a more conservative search strategy. The other two parameters, η and α were not significant predictors of either performance metric.

**Figure 4:**
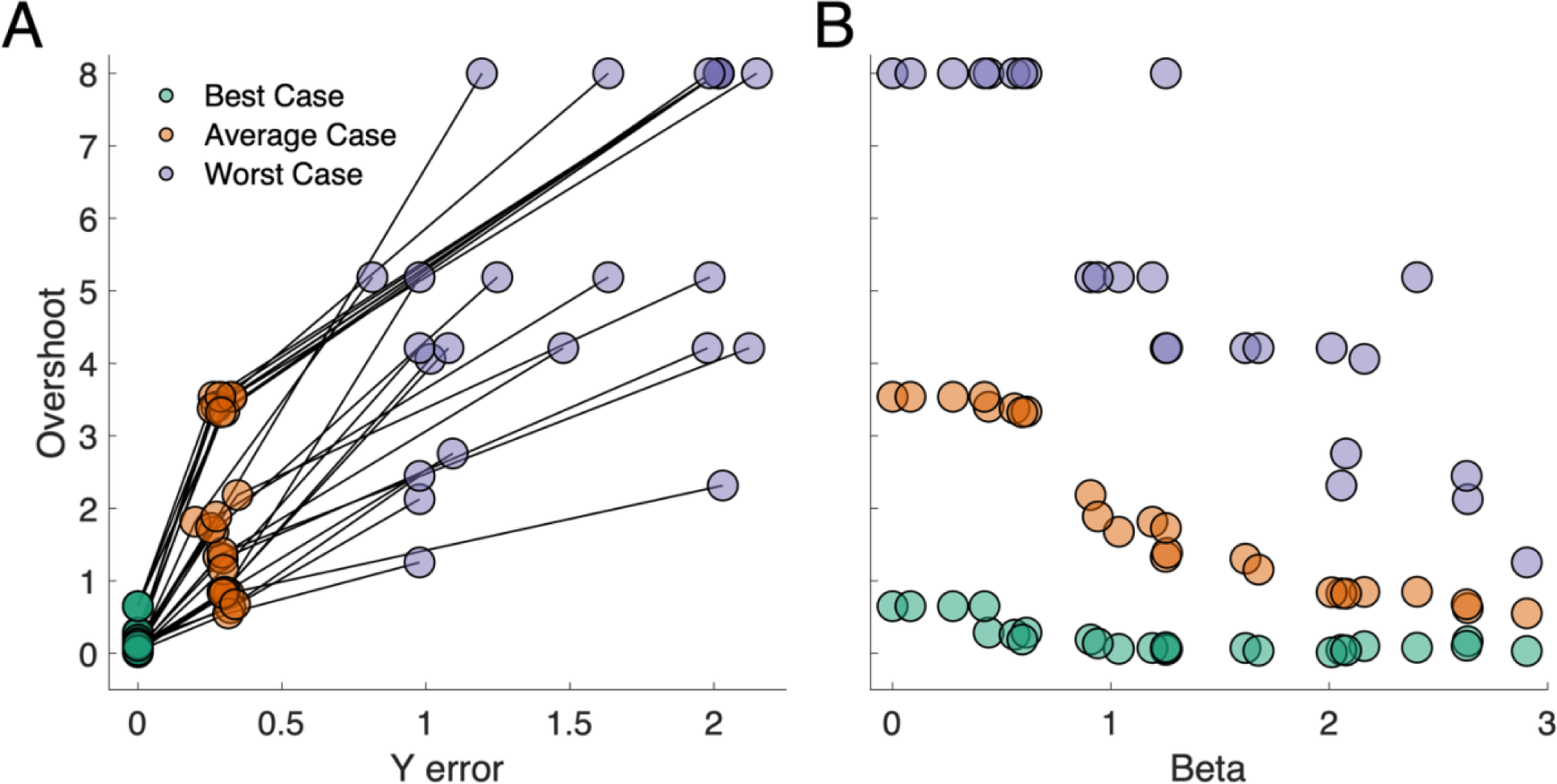
SAFE-OPT performance for *in silico* configuration parameter sweep. (A) The performance of each combination of configuration parameters in terms of final error and overshoot is represented by green, orange, and purple circles connected with black lines. (B) The overshoot of each combination of configuration parameters as a function of β.

### 3.4 *In vivo* validation

To validate our simulation experiments, we compared the performance of standard Bayesian optimization and SAFE-OPT *in vivo*. For SAFE-OPT, the η, β, and α configuration parameters that provided the lowest averaged final error and overshoot during the *in silico* prototyping were used when deployed in living subjects. For the standard Bayesian optimization algorithm, η was set to 0.4 to be consistent with previous DS optimization experiments.^26^

#### 3.4.1 Bayesian optimization for DA maximization

In subject S5 the stimulation amplitude parameter space was restricted to ±0-5 V. After the first three burn-in samples at ±0, 0.5, and 1.0 V the algorithm immediately selected the stimulation amplitude of ±4.0 V, followed by limited samples at ±5.0 V before it ultimately converged to an estimated optimal stimulation amplitude of ±3.95 V (**Figure 5A**). In the validation phase, the DS at ±4.0 V was not significantly different from the DS induced by sham stimulation, and the DA at ±4.0 V was higher than at sham (DA=0, by construction; **Figure 6A**). In subject S6, when the algorithm selected a stimulation amplitude of ±4.0 V, the stimulation induced a Racine level 5 behavioral seizure with rearing and falling. The fifth stimulation amplitude sampled by the algorithm, ±1.85 V also induced a seizure. Ultimately the algorithm converged onto an estimated optimal of ±0.7 V. This amplitude was validated against sham and ±1.5 V, which respectively produced a lower DA and DS.

**Figure 5:**
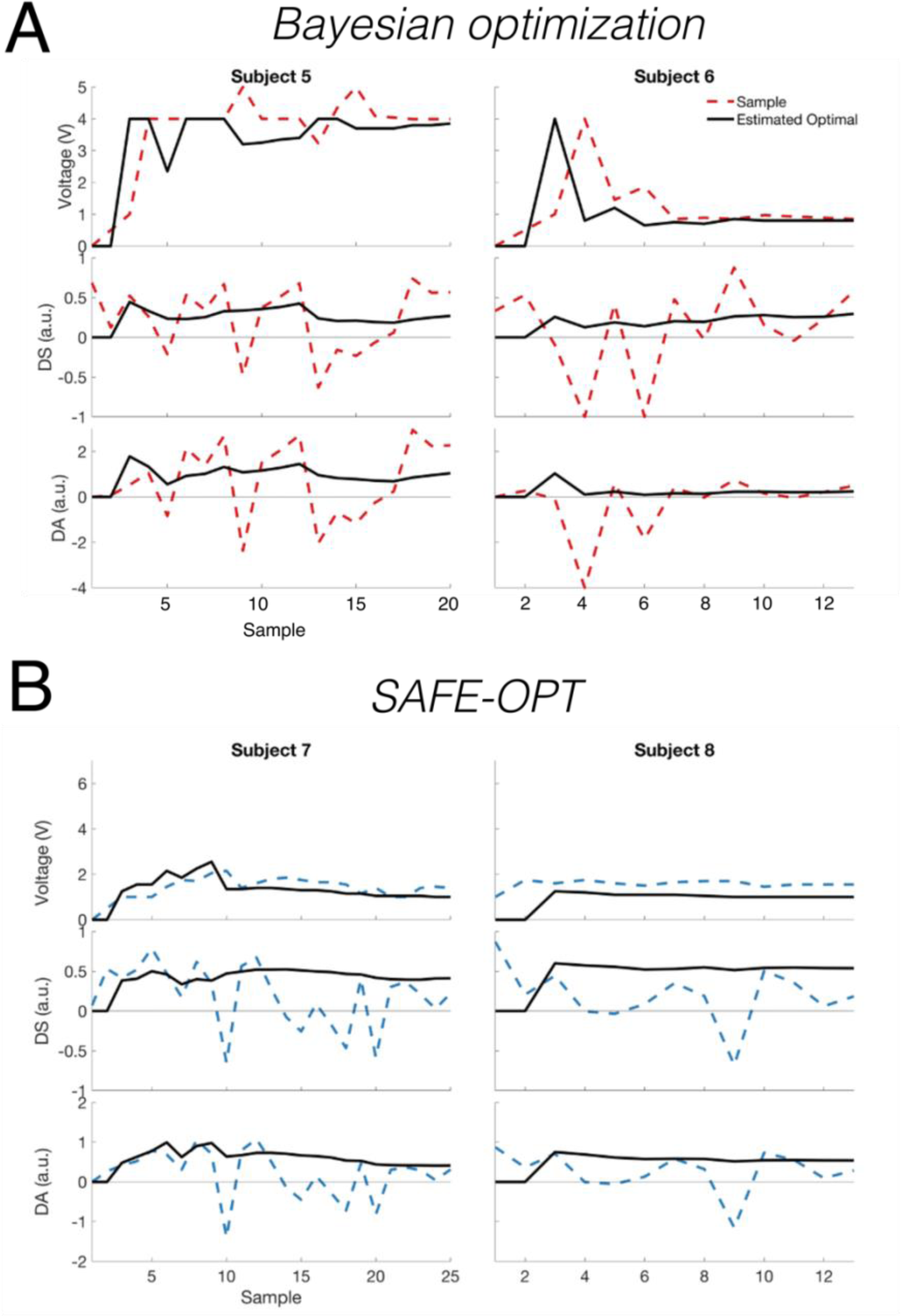
Search trajectories of Bayesian optimization and SAFE-OPT during DA maximization *in vivo*. Each column shows the series of input values (voltage), the safety function (DS), and the objective function (DA) selected by Bayesian optimization in A) and SAFE-OPT in B) at each iteration of a single optimization sequence. The dashed line indicates the data sample tested/collected at each iteration, while the thick black line indicates the optimal stimulation amplitude estimated by the algorithm and resulting DS and DA, according to the maximum expectation of the surrogate model.

**Figure 6:**
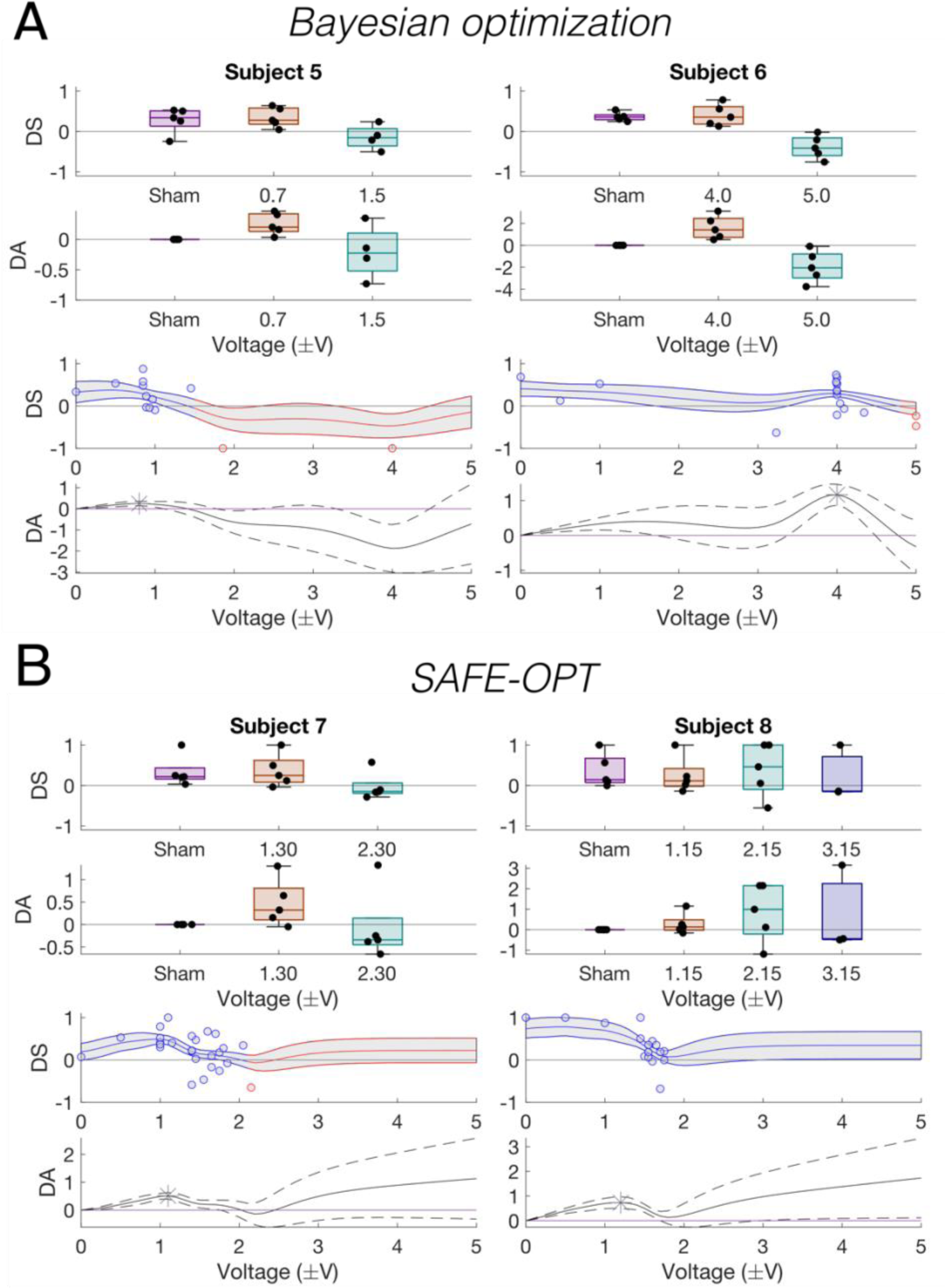
Bayesian optimization and SAFE-OPT validation results for maximizing discrimination area *in vivo*. Top: the DS and DA during sham, optimal, and control stimulation amplitudes. Each circle represents a single trial, box plots indicate range and quartiles, and the gray line indicates the mean value for each condition. Bottom: subject-specific ground-truth GP models of the expected DS and DA with confidence intervals using all data collected in this study. Blue/red line indicates the respective safe/unsafe regions of the parameter space according to the expected DS, and blue/red circles indicate safe/unsafe samples collected during optimization. Gray patch: 95% confidence interval estimated from the GP. Asterisk: optimum of the DA objective function.

#### 3.4.2 SAFE-OPT for DA maximization

The SAFE-OPT algorithm designed in section 3.3 was then deployed in *in vivo* in two additional subjects. In both subjects, SAFE-OPT employed a much more conservative strategy when choosing stimulation amplitude values, with ±2.1 V as the highest amplitude sampled (**Figure 5B**). In the validation phase for subject 7, the control stimulation at ±2.3 V produced a negative DS, while for subject 8, the initial control point at ±2.1 V did not cause a memory deficit. However, an additional control stimulation at ±3.15 V did cause a slight decrease in performance. These findings indicate that the SAFE-OPT algorithm was more successful in avoiding unsafe regions of the parameter space, but may have slightly underestimated the true safety boundary in one subject given the noisy nature of the behaviorally-measured objective function. There was an overall significant difference in DS measured for control point stimulation settings (p = 0.0013; Friedman test; n=4 subjects, 5 settings per subject) but not between sham and optimized stimulation values (p = 1.0; post-hoc multiple comparison test), indicating a successful optimization result for either approach.

## 4. Discussion

### 4.1 Summary

In this study we designed and characterized an algorithmic approach for safely optimizing brain stimulation. We demonstrated how the SAFE-OPT algorithm can efficiently maximize a noisy objective function based on a memory task, while simultaneously avoiding regions of the parameter space that may disrupt memory. In contrast, standard Bayesian optimization sampled stimulation settings that harmed task performance and caused seizures before arriving at the optimum.

### 4.2 Interpretation of results

The most evident challenge of this neuromodulation optimization problem was the magnitude of the noise – the sample-to-sample variability – of the behavioral task. Across the models generated from the eight animals in this study, the average contrast-to-noise ratio (CNR) was below 0.5, while in other studies^11, 30^, the CNR of the objective function could be up to an order of magnitude larger. Bayesian optimization was successful in optimizing the stimulation amplitude despite high noise in the objective function and the low number of task DS score samples that could be collected (1/day). While the standard Bayesian optimization approach successfully identified optimal stimulation settings, the exploration mechanism of the algorithm led it to test the extremes of the parameter space, which ultimately resulted in side-effects for one of the subjects.

In contrast, the SAFE-OPT approach was able to slowly broaden the search space, avoiding extremes of the parameter space unless necessary to find the optimum of the objective function. Given the noisy data, hyperpriors were also useful to prevent premature convergence or overshoot. While we evaluated the effect of three different configuration parameters on the performance of the overall SAFE-OPT algorithm, only β, the parameter controlling the magnitude of the safety constraint, had a substantial effect on performance. By increasing β, we could limit how much the stimulation amplitudes would overshoot the optimum with little effect on the other performance metric, final error.

### 4.3 Related work

Data-driven optimization approaches have been gaining traction for tuning brain stimulation in both clinical and pre-clinical studies. Bayesian optimization has been applied *in silico* for optimizing tACS stimulation,^16^ evoking movement with peripheral nerve stimulation for neuroprosthetic applications,^31^ and for optimizing STN stimulation based on evoked biomarkers that provide multiple, mutually competitive objectives.^30, 32^ Additionally, it has been deployed for modulating hippocampal oscillations through the adjustment of optogenetic stimulation parameters.^33, 34^

Two prior studies have explored the incorporation of safety constraints into Bayesian optimization for neural engineering applications, including accounting for patient side-effects during DBS programming for tremor relief^35^ and a computational exploration of design parameters that influence safe Bayesian optimization performance in a multidimensional input space.^36^ In this study, we demonstrate the successful application of safe Bayesian optimization in a rodent *in vivo* model, as well as a robust computational design methodology that can be replicated to design SAFE-OPT for best performance in future applications.

### 4.4 Future work

While we explored one possible implementation of Bayesian optimization with learnable safety constraints, safe optimization is a nascent field with many emerging advances and novel implementations.^37^ In particular, recent safe optimization approaches can leverage surrogate models more appropriate for high-dimensional parameter spaces to accommodate more complex stimulation parameter spaces.^38^ The mechanisms for learnable safety constraints can also be incorporated into multi-objective and state-dependent optimization problems. Safe optimization could also be applied further to safely guide the use of novel stimulation and recording paradigms.^39–41^ Ultimately, all of these approaches will be critical for developing safe and effective automatic optimization approaches for neuromodulation.

### 4.5 Limitations of the study

The results of this study should be interpreted based on the overall goal of designing a safe optimization algorithm using a proof-of-concept model system. While asynchronous distributed stimulation is a potential therapeutic option for drug-resistant temporal lobe epilepsy, and it will be crucial to consider any memory side-effects, this study was not designed to provide insight into the clinical or cognitive effects of the therapy. The algorithm designed for safely maximizing DA is specific to this optimization problem. However, these results show that the design framework employed in this study can be used to design problem-specific safe optimization algorithms that can achieve reliable performance for even very noisy behavior-based objective functions.

## 5. Conclusions

Bayesian optimization with learnable safety constraints can safely and efficiently identify stimulation settings that maximize a noisy behavior-based objective function while avoiding regions of the parameter space that may be unsafe.

## Author Contributions

M.J.C. developed the optimization framework, implemented the optimization experimental platform, conducted simulation experiments, and prepared the manuscript. E.R.C. performed experiments, data analysis, and prepared the manuscript. M.G., M. S., A.K. and T.E.G. conducted *in vivo* experiments and aided manuscript preparation. R.E.G. aided in project design, supervised experiments and helped prepare the manuscript.

